# Task prioritization modulates low frequency EEG dynamics reflecting proactive cognitive control

**DOI:** 10.1101/2022.05.04.490638

**Authors:** Nathalie Liegel, Daniel Schneider, Edmund Wascher, Stefan Arnau

## Abstract

Most neuroscientific studies investigating mental effort apply unspecific effort allocation paradigms. In contrast, the present EEG study targets specific effort allocation during task prioritization.

Twenty-eight participants performed a cued number classification task during the retention interval of a working memory task including retrospective cues. One of two possible number classifications was done per trial. Each trial started with a cue indicating which of the two tasks would be more important in the upcoming trial. Subjects were told to engage in both tasks, but to concentrate on the important one. Feedback given at the end of each trial was calculated based on task performance, with scores obtained from the relevant task being tripled.

Participants performed significantly better in either task when it was important compared to when not.Task prioritization modulates theta, alpha and beta oscillations, predominantly during task preparation. Multivariate pattern analysis revealed that the exact type of the two possible number classifications was decodable, however, decoding accuracy did not depend on task importance. Hemispheric alpha power asymmetries indicating attentional orienting between working memory representations also did not depend on task importance. The findings suggest that task prioritization primarily affects proactive cognitive control on a superordinate level.

## Introduction

The ability to prioritize the most important task, or the most important specific aspect of a complex task, is crucial to maintain a sufficient level of performance when dealing with a complex task or a multi-tasking scenario. It is well established that task performance does not only depend on task demands and processing capacities, but also on motivational factors ^1^. In this context, mental effort is the force that bridges the gap between being able to perform well in a task and indeed performing well ^1,2^. According to a theory by Shenhav and colleagues ^3^, the expenditure of effort is regulated by a task’s expected value of control (EVC, task importance). The EVC is conceptualized as the expected reward associated with successful task performance (further referred to as “reward”) minus its cost, i.e., the effort needed to solve the task. It could be shown that increasing a task’s importance, either by increasing its perceived outcome or by decreasing its perceived costs, positively affects task performance ^1,4,5^. In the present EEG study, we address the question of how task prioritization, as a special case of EVC manipulation, affects the EEG.

Recent neuroscientific studies provide evidence that the importance of a task affects various aspects of attentional and cognitive control ^4,6,7^. As attention and cognitive control often have overlapping definitions ^8^, we here use the term “attentional control” and “cognitive control” interchangeably in order to describe processes that keep our thoughts and actions aligned with our current goals and our goals updated ^4,8,9^. Most of the neuroscientific studies manipulating task importance use monetary incentives. A paradigm that is often used is the monetary incentive delay (MID) task ^10^. In this paradigm, a reward-cue prior to the actual target stimulus indicates how much monetary reward can be earned with successful task performance. At the end of each trial, a feedback screen reveals the earned reward.

Using this paradigm, fMRI studies have found that not just dopaminergic areas like the ventral striatum exhibit a stronger activation in high versus low reward conditions, but also frontal and parietal regions, which have been associated with task preparation and cognitive control. The MID paradigm has also been deployed in EEG studies. Several studies observed significant reward-related effects in response to the target stimulus. An enhanced parietal P3 ^11,12^, a stronger frontal N2 ^12^, as well as a more pronounced N2 posterior contralateral (N2pc) as an indicator for the selective orienting of attention ^13^ in the high compared to the low reward condition could be observed. Furthermore, reward-related effects were not only be found during actual task-processing, but also already during the cue-target interval. Here, a stronger desynchronization of alpha activity at central recording sites in the high versus low reward condition was observed in a visual search ^13^ and a Stroop task ^12^. High reward was also found to be related to enhanced amplitudes of the frontal CNV ^11,12,14^ during the cue-target interval, as well as to an increased parietal P3 ^14^. Taken together, these results indicate that task importance affects EEG correlates of cognitive control, as well as correlates of the allocation of cognitive resources.

The fact that task importance not only affected task-processing during the target-response interval, but also preparatory processes during the cue-target interval shows that proactive cognitive control mechanisms are modulated by the EVC. According to the dual mechanisms of control framework ^15^, proactive control refers to a processing mode during which task goals are actively maintained, whereas reactive control refers to a late correction mechanism which triggers the maintenance of goals when needed. The results from studies explicitly investigating the influence of reward on pro-versus reactive control using the AX-CPT, a variant of the continuous performance task, indicate that unspecific effort allocation predominantly modulates proactive rather than reactive control ^16–20^.

Another important aspect of the question how modulations of task importance affect cognitive processes is which levels of task processing are affected. With respect to how task prioritization is implemented, it is of interest to disentangle the contributions of cognitive control adaptations on a superordinate level on the one side, and on more specific *within task* level on the other side. The first refers to attentional selection differing one task from another task or differing task-relevant processes from task-irrelevant processes. The latter refers to attentional selection within a task, e.g., between two task stimuli. Behavioral studies show that task prioritization affects attentional resource allocation between tasks ^21–24^. However, to our knowledge it has not yet been investigated if task prioritization, i.e., specific effort allocation, also affects within-task attentional processes.

This issue has, however, been addressed in the context of unspecific effort allocation. A promising approach to investigate the influence of task importance on within-task processes is to decode the EEG signal using multivariate pattern analysis ^25,26^. Using a combination of a task switch and a MID paradigm in which the participants had to engage in two different classification tasks, Etzel et al. ^27^ decoded the classification tasks from fMRI data, for rewarded and unrewarded trials separately. Hall-McMaster et al. ^28^ applied a similar analysis using EEG data. In both studies, the decoding analysis revealed that activity patterns associated with the classification tasks were more distinct from each other in rewarded than in unrewarded trials. The EEG study furthermore revealed that also task stimuli and response sides were more distinct from each other in case of a high task importance. Hence, in these studies, task importance affected differences at the within-task level.

Another way to isolate within-task processes is to make use of lateralized stimuli. In specific working memory tasks, a retrospective cue (retro-cue) enables the subjects to shift their attention to only one of two laterally presented items held in working memory. Previous research found that such shifts of spatial attention are reflected by hemispheric alpha power asymmetries ^29,30^. Task importance has repeatedly been observed to modulate spatial attention reflected by lateralized ERP measures like the N2pc and the CDA ^13,31,32^.

The goal of the present study is to investigate task prioritization in a complex task. To our knowledge, there is no study investigating neural correlates of task prioritization, i.e., of the task-specific manipulation of the EVC in a way that some task-relevant processes are more important than others. By using the EEG, we want to address the question in how far pro-versus reactive control modes on the one side, and superordinate versus within-task control processes on the other side are affected by modulations of task prioritization. We use a nested design in which a working memory task is interrupted by a cued number classification task. For the working memory task, the subjects must remember two laterally presented stimuli. A retro-cue appearing after the completion of the number classification task then indicates which stimulus needs to be recalled. In the cued number classification task, the cue indicates whether the participants must classify a single digit number as either bigger or smaller than five, or as odd or even. For each trial, one of the two tasks is cued as being more important. A combined feedback score is presented at the end of the trial, with the score obtained from the respective important task being weighted three times as strong as the scores obtained in the less important task. In contrast to previous studies investigating task importance, the general amount of reward that can be earned is the same for each trial. Therefore, the reward cue at the beginning of a trial does indicate *when* instead of *if* effort needs to be allocated to maximize the reward.

Behavioral task prioritization research has shown that attentional resources can be willfully allocated to certain aspects of a task or to one of two concurrent tasks ^21–24^. We therefore expect the participants to perform more accurate in the working memory task in trials in which the working memory task is more important, and to perform better in the number classification task in trials in which the number classification task is more important.

EEG data were analyzed in two different ways, exploratory and task specific. First, we performed an exploratory analysis on the time-frequency decomposed signal to investigate how task prioritization influences attentional control in general and pro versus reactive control in specific. Recent research suggests that different ways to manipulate task importance affect attentional and cognitive control similarly ^7,33–35^, We therefore expect task prioritization to not be an exception and to primarily affect low-frequency oscillations which are known to reflect the exertion of cognitive control ^36–38^. In particular the results from cued reward (MID) paradigms suggest that task importance mainly affects proactive cognitive control ^20,28,34,39–42^. Since the position of the important task within each trial is known at the beginning of each trial in the present study, we expect task importance to influence oscillatory activity during preparatory time intervals: not only during the relevance cue, but also during other time intervals preceding the start of a task, e.g., during the number classification task cue.

Second, we performed task-specific analyses to investigate whether prioritization rather affects superordinate or within-task processes. For the number classification task, we used multivariate pattern analysis to measure the distinguishability of the two different number classification rules in the signal. To understand if task prioritization affects how strongly the different task rules are represented in the signal, we decoded the classification type for number classification task important and working memory task important separately. A higher decoding accuracy in trials in which the number classification task is more important than in trials in which it is less important would indicate that task prioritization influence attentional control on a within-task level. As an index of within-task control in the working memory task, the spatial shift of attention within working memory in response to the retro-cue was targeted, which should be reflected in topographical asymmetries of alpha power. Here, we can conclude that task prioritization affects within-task attentional control if the strength of alpha power asymmetry is found to depend on task importance.

## Results

### Behavioral data

Subjects performed better in either task when it was important compared to when not.

In the number classification task, feedback scores depended on both reaction time and accuracy of answers. Both measures provided therefore an index of task performance. Participants answered significantly faster when the number classification task was important (*M* = 537.00 ms, *SD* = 114.13 ms) than when the working memory task was important (*M* = 562.86 ms, *SD* = 119.61 ms; *t*(27)=5.42,*p* < .01, *η* = 0.50, *d* = 1.02). No significant difference between working memory task important (*M* = 8.66%, *SD* = 6.38%) and number classification task important trials (*M* = 8.56%, *SD* = 5.62%) was found regarding accuracies (*t*(27)=.27, *p* = .79).

In the working memory task, participants were instructed to answer as accurately as possible and only accuracies, not reaction times contributed to feedback scores. Accuracies were measured as the degree deviation between the response orientation given by the participant and the to-be-remembered orientation presented earlier in the trial. Participants answered significantly slower, probably considering their answer more thoroughly, in the working memory task important condition (*M* = 1368.91 ms, *SD* = 300.92 ms) than in the number classification task important condition (*M* = 1292.68 ms, *SD* = 267.38 ms; *t*(27)=5.01,*p* < .01, *η* = .46, *d* = .95). For degree deviations the pattern was reverse, meaning that degree deviations were significantly smaller in the working memory task important condition (*M* = 11.77°, *SD* = 4.07°) than in number classification task important condition (*M* = 13.48°, *SD* = 5.23°; *t*(27)=-3.80,*p* < 0.01, *η* = 0.32, *d* = 0.72).

### Mass-univariate analysis of time-frequency decomposed EEG data

A cluster-based permutation test on decibel-normalized power values in the electrode × time × frequency space comparing the number classification task important and the working memory task important condition resulted in four significant clusters (see figure 2). Two clusters, one in the theta range (2 to 7 Hz) and one in the alpha/beta range (7 to 20 Hz), subsequently referred to as beta cluster, were significant (theta cluster: *p* < .01, beta cluster: *p* = 0.02) during and after relevance cue presentation (theta cluster: −815 to 850 ms relative to relevance cue; beta cluster −150 to 1025 ms relative to relevance cue). Oscillatory activity was significantly stronger in number classification task important trials than in working memory task important trials in both clusters (see figure 2D). The beta cluster (cluster 1 in figure 2) showed strongest differences between both experimental conditions at left occipito-parietal, frontal, and right parietal electrode sites (see figure 2A), the theta cluster (cluster 2 in figure 2) at right parietal sites (see figure 2B). In between the onset of the memory items and the start of the number classification task (significant at 2770 to 5660 ms relative to relevance cue onset), another cluster was significant (*p* = 0.02) in the theta frequency range (significant at 3 to 9 Hz). This cluster, like the theta cluster during relevance cue, showed stronger values for number classification task important trials and was most pronounced at right parietal electrode sites (see figure2C). Finally, a fourth cluster was significant (*p* < 0.01) in a broad frequency range from 6 to 15 Hz and from 5665 to 7830 ms relative to relevance cue, i.e., during number classification task performance. Differences between experimental conditions in this cluster were most pronounced at parieto-temporal electrodes sites (see figure 2E) and in the alpha frequency range (see figure 2D). This was the only cluster showing stronger activity for working memory task important than for number classification task important trials. All cluster were significant at nearly all electrodes: The early theta cluster (cluster 2 in figure 2) was significant at all channels except TP9, TP10, PO9, PO10, FT9, FT10, C5, the other three cluster were significant at all channels except TP9, TP10, PO9, PO10, FT9, FT10.

**Figure 1.**
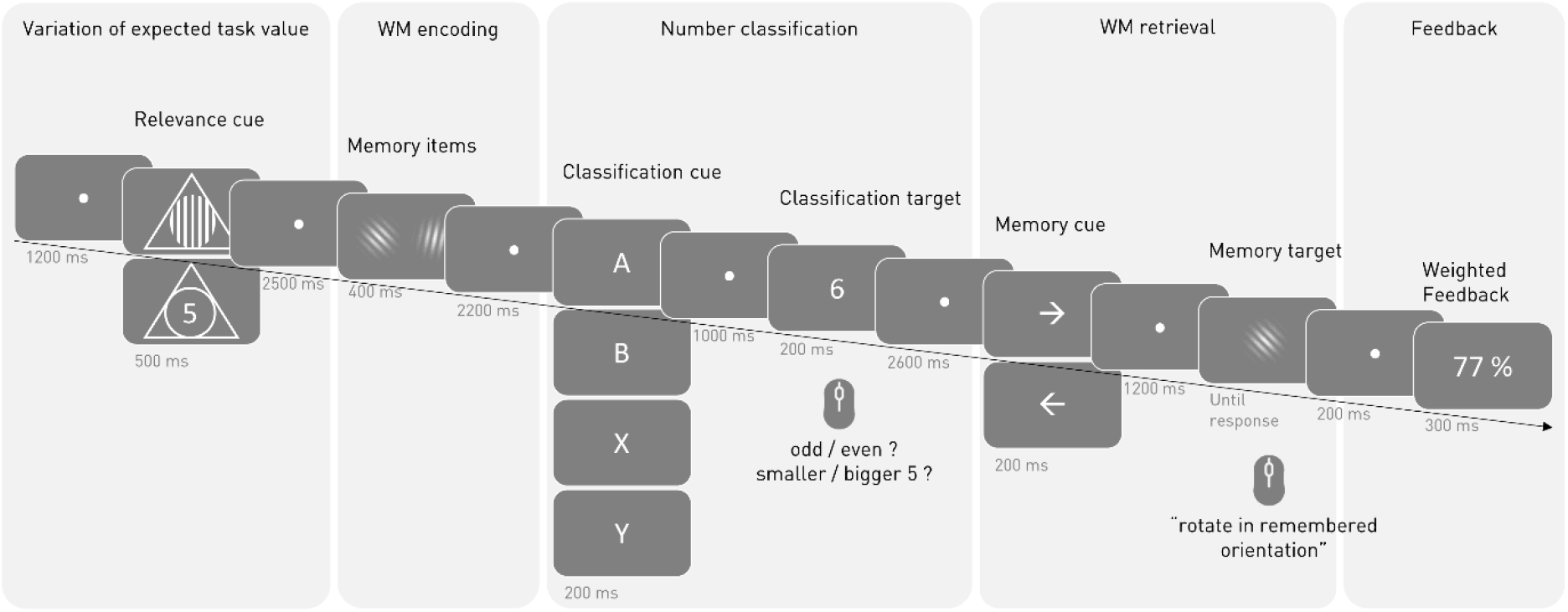
Schematic illustration of a trial’s sequence. Each trial consisted of five parts depicted here by five bright grey columns. The trial started with one of two possible relevance cues indicating which of two tasks would be more important and thus should be focused on during the upcoming trial. Then, participants performed a cued number classification in the retention interval of a working memory task. In the number classification task, participants classified the classification target (one of the digits 1,2,3,4,6,7,8,9) via left or right button press. The classification cue indicated which of two classifications should be done. In the working memory task, participants used their mouse to rotate a Gabor patch with random orientation (the memory target) in a previously remembered orientation of a memory item. In each trial, two memory items are shown, one on the left and one on the right side of the screen. The memory cue pointed to the side of the memory item which should be used in the memory task. At the end of each trial, participants perceived feedback based on their performance in both tasks. Performance in the more important task was weighted three times stronger than performance in the less important task. The bright grey numbers under the schematic screens indicate stimuli’s duration on screen. WM = working memory.

**Figure 2.**
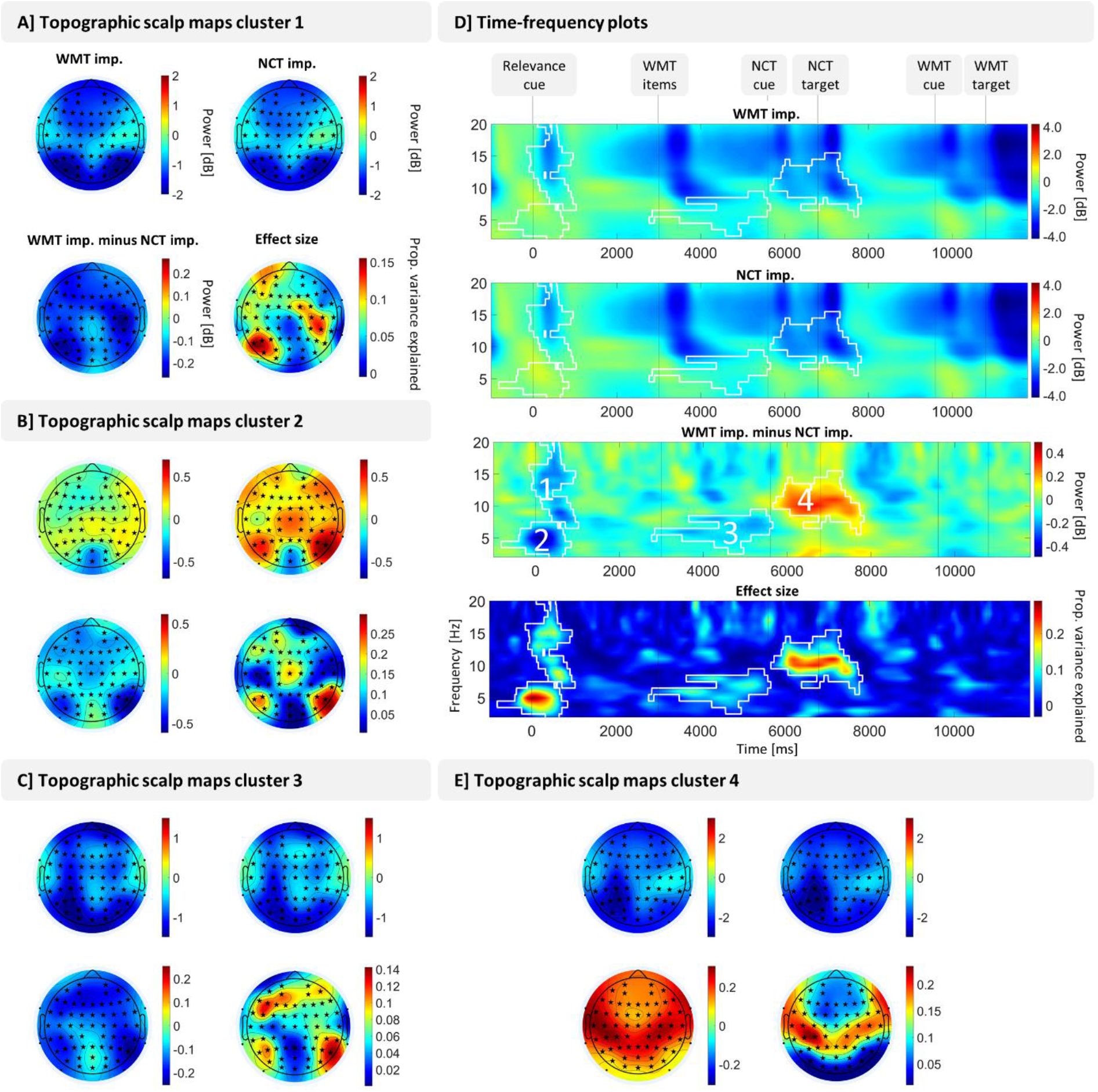
Result of univariate power analysis. A cluster-based permutation test in the electrode × times × frequency space comparing power in number classification task important (NCT imp.) trials and working memory task important (WMT imp.) trials revealed four significant clusters. Panel A, B, C and E show topographic scalp maps averaged over significant time-frequency points of the respective cluster. Significant channels are marked with a black asterisk, not significant channel with a black dot. D shows time-frequency plots with power values averaged over all channels (as the four clusters were all significant at nearly all channels). Contour lines frame significant time-frequency points and the numbers label the cluster from 1-4. Effect sizes are adjusted partial eta squared. Power values are decibel-normalized with a condition-unspecific baseline ranging from 700 to 200 ms prior to the onset of the relevance cue.

### Multivariate pattern analysis

We decoded number classification type (smaller/bigger five vs. odd /even) based on ERP values and alpha power values (8-15 Hz) for a time interval covering the whole trial (from relevance cue onset until one second after working memory target onset). We were especially interested in alpha power as mass-univariate analysis of time-frequency decomposed EEG data revealed significant changes between experimental conditions during number classification task performance in this frequency range. Therefore, task importance obviously has an effect on alpha power. To better understand what neural mechanisms are reflected by alpha activity, decoding analysis aimed to reveal if task importance also affects how distinguishable classification type task sets are from each other within the alpha range.

Figure 3 shows decoding accuracies averaged over participants. In both cases, ERP decoding (see figure 3A) and power decoding (see figure 3B), accuracies slightly fluctuate around the chance level at 50% and peak after the onset of the number classification task cue and the number classification task target. Approximately one second after number classification task target onset, curves bottom out at chance level again.

**Figure 3.**
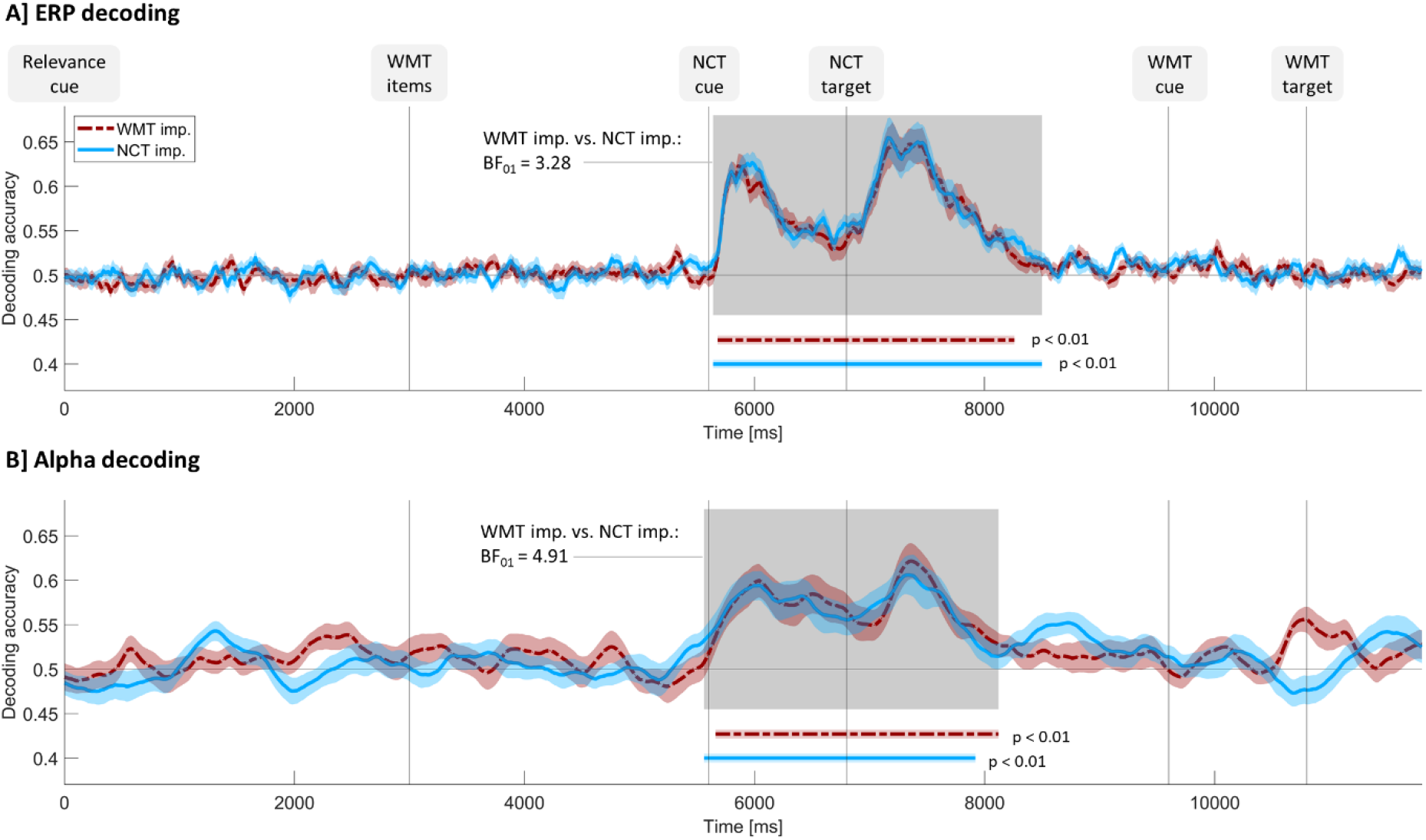
Decoding accuracies over time for ERP (panel A) and alpha power (panel B) for both experimental conditions separately. The horizontal line at 0.5 indicates chance level. Vertical lines indicate stimuli onset times. Shaded curve surroundings indicate between -subject standard error of the mean. Vertical colored lines mark significant time points according to cluster-based permutation tests over time. The grey shading indicates time points where at least one experimental condition differs significantly from zero according to those tests. BF_01_ = reverse Bayes factor calculated for decoding accuracies averaged over the time points marked by the grey shading; WMT = working memory task; NCT = number classification task; imp. = important.

Cluster-based permutation test against chance level revealed a cluster of significant time points during number classification performance for both experimental conditions and both decoding types (ERP decoding, working memory task important: *p* < .01, 5680 to 8260 ms; number classification task important: *p* < .01, 5640 to 8500 ms; alpha power decoding, working memory task important: *p* < .01, 5660 to 8120 ms; number classification task important: *p* < .01, 5560 to 7920 ms). As expected for an unbiased classifier, confusion matrices revealed similar decoding accuracies for both classes (mean decoding accuracies averaged over significant time points: ERP decoding, working memory task important, odd / even trials: .58, bigger / smaller five trials: .58; ERP decoding, number classification task important, odd / even trials: .57, bigger / smaller five trials: .58; alpha power decoding, working memory task important, odd / even trials: .57, bigger / smaller five trials: .57; alpha power decoding, number classification task important, odd / even trials: .57, bigger / smaller five trials: .57). Cluster-based permutation tests which compared number classification task important trials and working memory task important trials resulted in no significant clusters, neither for ERP decoding, nor for alpha power decoding.

*T* tests combined with Bayesian analysis revealed the same pattern of results: Bayesian one-sample *t* tests comparing chance level against decoding accuracy averaged over – according to cluster-based permutation tests – significant time points revealed that decoding accuracy is significantly above zero in all cases (ERP decoding, working memory task important: average over 5680 to 8260 ms,*M* = .58, *SD* = .05; *t*(27)=60.26,*p* < .01, *η* = .99, *d* = 11.39, BF_10_ = 5.99 * e^26^; ERP decoding, number classification task important: average over 5640 to 8500 ms; *M* = .58, *SD* = .05; *t*(27)=64.18,*p* < .01, *η* = 0.99, *d* = 12.13, BF_10_ = 3.04 * e^27^; alpha power decoding, working memory task important: average over 5660 to 8120 ms, *M* = .57, *SD* = .06; *t*(27)=50.22, *p* < .01, *η* = 0.99, *d* = 0.49, BF_10_ = 5.47 * e^24^; alpha power decoding, number classification task important: average over 5560 to 7920 ms,*M* = .57, *SD* = .06; *t*(27)=51.09,*p* < .01, *η* = .99, *d* = 9.66, BF_10_ = 8.51 * e^24^).

Bayesian paired *t* tests between experimental conditions were computed for the averages over time points in which at least one experimental condition differed significantly from chance level (ERP decoding: 5640 to 8500 ms, alpha power decoding: 5560 to 8120 ms), resulting in no significant differences (ERP decoding: working memory task important: *M* = .58, *SD* = .05; number classification task important: *M* = .58, *SD* = .05; *t*(27) = .99, *p* = .35; alpha power decoding: working memory task important: *M* = .57, *SD* = .06; number classification task important: *M* = .57, *SD* = .06; *t*(27)=.19, *p* = .85). The reverse Bayes factors of 3.28 for ERP decoding and 4.91 for alpha power decoding indicate substantial evidence in favor of the null hypothesis for both decoding types.

### Alpha power asymmetry

Hemispheric power asymmetries relative to the trial’s working memory retro-cue direction are shown in figure 4. More exactly, we quantified asymmetries by subtracting power at the hemisphere contralateral to the cued memory item from power of the hemisphere ipsilateral to the cued memory item. To normalize power values, this was then divided by total power across both hemispheres. This resulted in a lateralization index which is zero if there is no power difference between both hemispheres, positive in case contralateral alpha activity is weaker in the hemisphere in which visual processing of the memory item was done than in the other hemisphere and negative if it is the other way round.

**Figure 4.**
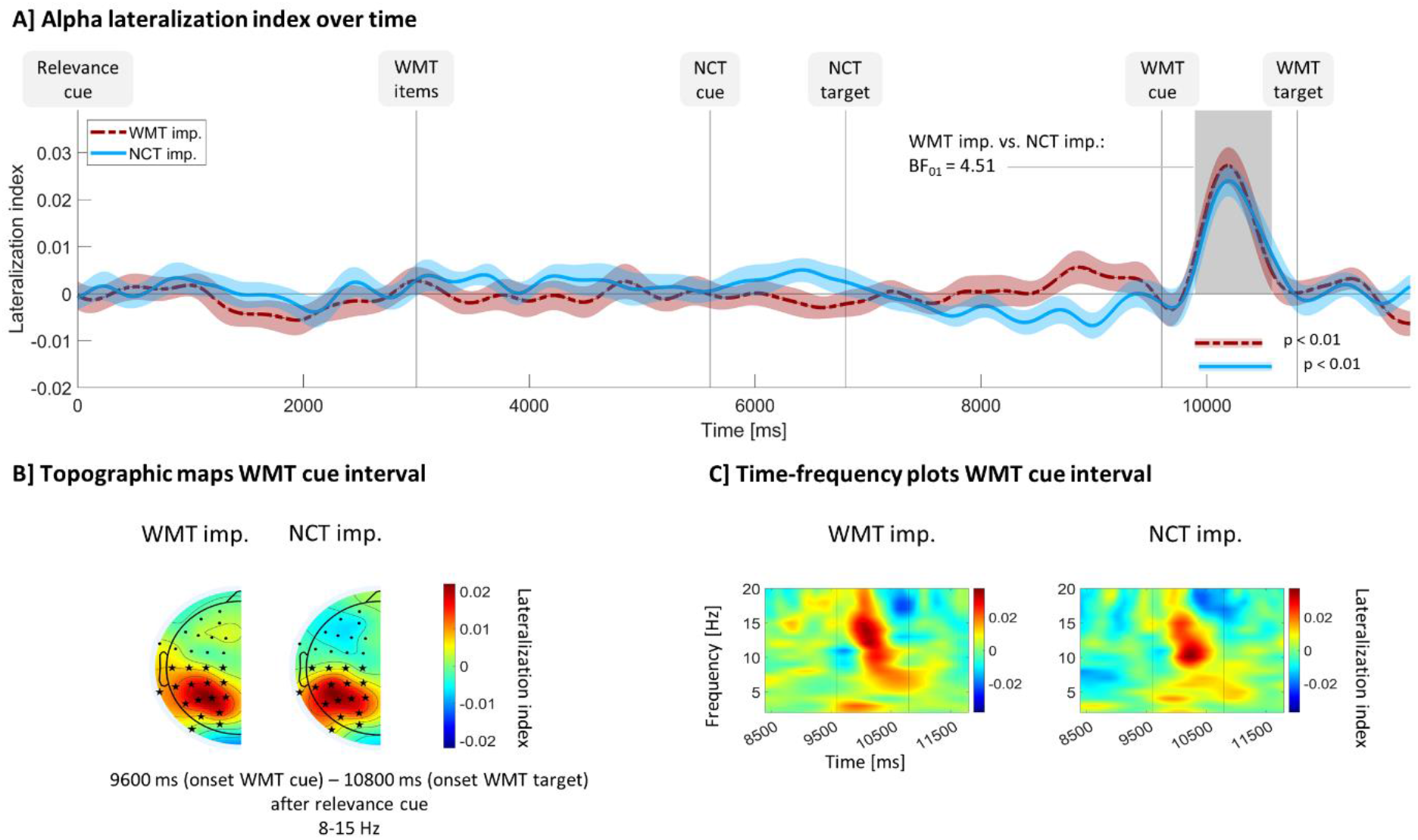
Alpha power asymmetry relative to WMT cue direction. All panels show lateralization index values, i.e., raw power at the hemisphere ipsilateral to the side the working memory task cue points to minus raw power contralateral to this side divided by raw power across both hemispheres. Index values above than zero are thought to reflect greater attention toward the cued memory item relative to the un-cued item. A) Time course of lateralization index averaged over 8-15 Hz and occipito-parietal electrodes (see electrodes marked with an asterisk in panel B). Shaded curve surroundings indicate between-subject standard error of the mean. Horizontal lines mark stimuli onsets and vertical, colored lines mark significant time points according to cluster-based permutation tests over time. The grey shading indicates time points where at least one experimental condition differs significantly from zero according to those tests. BF01 = reverse Bayes factor calculated for lateralization indices averaged over the time points marked by the grey shading. B) Topographic scalp maps of lateralization index averaged over 8-15 Hz and a time interval starting with working memory task cue onset and ending with working memory task target onset. Electrodes that were used for statistical analysis are marked with a black asterisk; other electrodes are marked with a black dot. As lateralization indices measure the hemispheric difference, scalp maps are per definition symmetric and only one half is displayed here. C) Frequency distribution of lateralization index around the working memory task cue onset. Horizontal lines indicate working memory task cue and working memory task target onset. Times are relative to relevance cue onset. WMT = working memory task; NCT = number classification task; imp. = important.

The lateralization index was strongest at around 10 Hz in number classification task important trials and at around 12-15 Hz in working memory task important trials (see figure 4C). It was most pronounced for both experimental conditions at occipito-parietal electrode sites (see figure 4B). Figure 4A shows the time course across the trial of lateralization indices, averaged over 8-15 Hz and occipito-parietal electrodes, for both experimental conditions. Index values remain close to zero except for a time interval in between the onsets of the working memory retro-cue and the working memory task target in which they are clearly bigger than zero in both trial types. A cluster-based permutation test over time revealed that lateralization indices differed significantly from zero during this interval for both, working memory task important (*p* < .01, 9895 to 10490 ms relative to relevance cue onset), and number classification task important condition (*p* < .01, 9930 to 10575 ms relative to relevance cue onset). Another cluster-based permutation test comparing both conditions resulted in no significant clusters throughout the whole trial.

To better understand these effects, *t* tests and Bayes factors were additionally computed for specific time intervals of interest: A one-sample *t* test comparing number classification task important lateralization indices with zero was computed on the average over those time points that were significant in the number classification task important vs. zero cluster-based permutation test, i.e., from 9930 to 10575 ms. This *t* test was significant (*M* = .02, *SD* = .02; *t*(27)=6.07,*p* < .01, *η* = .56, *d* = 1.15) and the corresponding Bayes factor of BF_10_ = 10430 indicates a strong effect in favor of the alternative hypothesis. An analogous *t* test on working memory task important lateralization indices (*M* = .02, *SD* = .02; *t*(27)=5.53,*p* < .01, *η* = .51, *d* = 1.04) averaged over time points in which working memory task important indices differed significantly from zero (9895 to 10490 ms) also resulted in a high Bayes factor (BF_10_ = 2833) indicating a strong effect in favor of the alternative hypothesis. A paired *t* test of working memory task important vs. number classification task important lateralization indices averaged over the time interval in which at least one condition showed significant effect according to cluster-based permutation tests (9895 to 10575 ms) showed no significant effects (number classification task important: *M* = .02, *SD* = .02; working memory task important: *M* = .02, *SD* = .02; *t*(27)=.47, *p* = .64). The Bayes factor of .22 (corresponding to a reverse Bayes factor BF01 = 4.51) indicates that data are 4.5 times more likely if there is in fact no difference between both experimental conditions, and thus, provides substantial evidence against the alternative hypothesis.

## Discussion

In the current study, we investigated neural correlates of mental effort using a dual task paradigm consisting of a working memory and a number classification task. In each trial, subjects were instructed to focus on one of two tasks. At the end of each trial, they received a combined feedback based on their performance in both tasks. Performance in the important task was weighted stronger than performance in the unimportant task. As expected, this instruction to adapt mental effort in a task-specific way resulted in significantly better behavioral performance in a task, when it was important than when it was unimportant ^21–24^. The EEG data was analyzed regarding three aspects: 1) Which changes in event-related oscillatory power can generally be observed? 2) Does the mental effort manipulation affect how distinguishable different number classification task types are represented in the signal? 3) Does the mental effort manipulation influence attentional orienting between working memory representations, reflected by alpha power asymmetry in response to a retro-cue? Question one addresses the aspect in how far task prioritization affects proactive vs. reactive control, question two and three in how far task prioritization affects cognitive control at a superordinate vs. a lower within-task level.

Question one was approached by an explorative cluster-based permutation test comparing the number classification task important with the working memory task important condition. Significant differences were identified during and after the cue indicating which of the two tasks would be more important. Oscillatory activity in a broad range of frequencies mainly over frontoparietal electrodes was stronger when the number classification task was important than when the working memory task was more important. Effect sizes were most pronounced for theta activity at five Hz at parietal electrodes. To interpret this effect, one has to consider that the relevance cue was different in both experimental conditions and was not counterbalanced across participants. Hence, early visual reward cue processing should be different for both experimental conditions and might result in measurable differences in the EEG. However, the very modest visual differences between both cues are unlikely to explain the effect that our analysis revealed: This effect extends over a time interval of nearly two seconds and over electrodes at nearly the whole scalp. Therefore, we interpret the observed differences to more likely reflect changes of higher-order preparatory processes due to task prioritization than to reflect changes in visual processing due to visual cue differences. Such preparatory processes might involve integration of the reward cue information in the current brain configuration and subsequent top-down biasing ^9^, strategic planning of effort allocation for the upcoming trial involving cost-benefit analyses in which the to be invested effort is weighed against the expected reward ^4,62^ and prospective memory activation as subjects have to plan *when* instead of *if* to invest effort. It is plausible that beta and theta oscillatory activity reflect those higher-order preparatory processes. Midfrontal theta is well-known to be involved in the process of flexibly updating the current brain configuration in response to environmental changes ^36^ and more specifically, pre-stimulus fronto-parietal theta has been interpreted to reflect goal updating ^63^ and top-down task preparation ^64,65^. In MEG ^66^ and EEG ^67^ studies in which a relevance cue indicates if an upcoming trial will be rewarded or not, theta and beta power modulations in response to the cue have been found. This suggests that the integration of reward information in the current task set can indeed be reflected by oscillations in the theta and beta range.

Another significant cluster was found in the time interval between the presentation of the working memory items and the start of the number classification task. In this time interval, event-related theta power throughout the whole scalp was significantly stronger in number classification task important condition than in working memory task important condition. The effect was strongest at frontoparietal electrodes. Regarding the working memory task, this contrasts with recent literature. Theta power is well-known to be related to encoding processes and to the maintenance of task-relevant information ^68^. However, theta power is usually increased in trials with a better performance ^38^, whereas in the current study, theta power is weaker in the condition associated with a better working memory performance. In addition, Kawasaki and Yamaguchi ^69^ found that high reward increased event-related theta power at frontal electrodes during the retention interval. As the result pattern is the opposite here, the observed modulations of theta power might reflect effort-related modulations of proactive cognitive control related to the number classification task. In line with this idea, frontoparietal ^70,71^ and frontal theta power ^72^ has been observed to be increased in response to a cue indicating a larger need for cognitive control in the upcoming trial compared to cues indicating that cognitive control will be needed to a lesser extent. Therefore, the increase in theta could reflect stronger goal updating and task preparation when the number classification task is more important compared to when the working memory task is more important.

Finally, a modulation of event-related frontoparietal alpha power was observed during the performance of the number classification task. Alpha power was reduced when the number classification task was important. This effect was strongest during the number classification task’s cue-target interval.

Overall, the observed modulations of event related spectral power changes related to the experimental manipulation occurred mainly in time intervals not directly related to task processing, that is not as an immediate response to imperative stimuli. These modulations are therefore likely to reflect proactive cognitive control. While the exploratory analysis in time-frequency space mainly revealed *when* during the trial the experimental manipulation affected task processing, a multivariate pattern analysis brought some insight regarding the question *on which level* effort affects task processing. In this analysis, we decoded from the EEG data whether an odd versus even or a smaller versus bigger five classification needed to be done in the number classification task, specifically addressing the question how strongly the difference in between both number classification task types is represented in the signal and if the effect of task prioritization gets through to lower-level, within-task processes of the number classification task or stays on a more general, superordinate level. Classifiers were trained for each time point throughout the trial using ERP data. We observed above-chance decoding accuracies for the time interval of the number classification task, indicating that the type of the number classification task is generally decodable from the EEG scalp topography. However, the observed classifier accuracy did not differ between experimental conditions, suggesting that task importance did not affect the distinguishability of the task representations in the EEG signal.

We did the same decoding analysis for alpha power topographies as, during the number classification task, reward seems to mainly have an influence in the alpha frequency range according to the cluster-based permutation test described above. A potential effect of task importance on number classification task type might be more pronounced when restricting the signal to the relevant frequency range. In fact, with alpha power, we obtained the same result than with ERP data: Classification type was decodable throughout the whole time interval during which the number classification task was done but task importance did not influence this decodability. This null finding helps to further interpret the significant difference in alpha activity between the number classification task important and working memory task important condition obtained in the cluster-based permutation test: The condition difference in alpha activity seems to not reflect condition differences at a within-task level, i.e., differences in rule-specific task preparation and task processing. It rather seems to reflect resource allocation on a more superordinate level.

However, it remains speculative to better define these superordinate processes underlying the alpha effect obtained in the cluster-based permutation test. Alpha power is thought to reflect attentional biasing ^73–76^. A decrease of scalp alpha is often observed when attention is directed to visual objects whereas an increase in scalp alpha appears when attention is directed to internal thoughts, mind-wandering and mental imagery ^38,77–79^. In accordance with this, we could explain the increase in alpha during the number classification task when the working memory task is important by both, an enhanced shielding of the to-be-remembered working memory representation and, by decreased attention to the number classification task. The first interpretation is supported by behavioral multitasking studies in which task prioritization has been found to lead to protection of the prioritized task against interference from a second task ^21,24,80^. The second interpretation is supported by previous studies revealing that a decrease of alpha power in the case of high task importance has been observed repeatedly for single tasks, hence in situations where no dual task shielding is necessary ^12,13,81^. Furthermore, as the alpha effect is most pronounced at parieto-temporal electrodes and is slightly stronger over the left hemisphere (i.e., contralateral to the response hand), the effect might to a large degree reflect pre-motor and motor preparation for the number classification task The most likely explanation is that in our study, processes related to the working memory and to the number classification task contribute to the alpha effect. This is supported by the observation that the effect is time-locked to the start of the number classification task, hence, to the start of the dual-task time period.

To sum up, although it remains a matter of debate if the condition difference in alpha activity reflects shielding of the working memory task or an attentional strengthening of the number classification task, we can conclude that the difference underlies a rather general, superordinate resource allocation process which does not generalize to a lower within-task level.

In a third analysis, we investigated such a within-task process in the working memory task. More specifically, we analyzed the hemispheric asymmetry of alpha activity in response to a retro-cue indicating which of two items hold in working memory should be remembered and which could be forgotten. Earlier research indicates that this asymmetry reflects a shift of attention toward the cued working memory representation ^29,30^. A significant lateralization of alpha power with higher power ipsilateral to the cued site was observed, but this effect was unaffected by task importance.

One possible explanation is that processes at within-task levels are generally not affected by task prioritization. This fits with the other results obtained in this study. However, there are other possible explanations: It is possible that the late time interval of the trial was not affected at all – on neither level – as earlier, proactive resource allocation was sufficient to efficiently allocate effort between tasks. This interpretation is in line with the finding that reward in working memory tasks often has its strongest influence during encoding and maintenance ^4,82,83^. To conclude, our finding is in line with the assumption that task prioritization mainly exerts its influence on proactive and superordinate attentional control processes. Future research might further specify how effort affects working memory processes in general and attentional orienting between working memory representations in specific.

## Conclusion

To sum up, in contrast to previous neuroscientific research in this field, we here manipulated mental effort in a task prioritization paradigm. Although we used a new manipulation of effort allocation, the goal of the current study was not to directly compare different types of task importance operationalizations as has been done before ^33,35,84^. Instead, we aim to extend the findings of previous studies to a broader context. Our results enable to understand neural mechanisms underlying task prioritization mainly regarding two aspects: First, the task involved several task cues allowing to isolate proactive cognitive control processes. Our results indicate that task prioritization mainly affects task preparation, i.e., proactive control. In this context, we show that effects of task importance on theta oscillations are not restricted to the time interval directly following reward cue processing as found in earlier studies ^66,67^. Instead, here, we found a modulation of theta activity that seems to reflect preparation of the upcoming task and is completely independent of reward cue processing. Second, we ask if task prioritization rather affects superordinate strategic processes or task-related activity. Results suggest that task prioritization rather affects general cognitive control at superordinate levels instead of within-task attentional selection.

In a nutshell, we found that task prioritization modulates on the one hand task preparation, reflected mainly in theta oscillatory activity, and on the other hand resource allocation during task processing, reflected in alpha oscillations. Hence, according to our results, preparative instructions how to direct one’s focus in dual task situations modulate strategic planning and attentional resource allocation across tasks rather than facilitating task processing itself.

## Methods

### Participants

Twenty-eight volunteers (9 men, 19 women) with a mean age of 23.13 years (*SD* = 3.64), with normal or corrected to normal vision and with no psychological or neurological diseases were included in the experiment. All of them were right-handed according to the Edinburgh Handedness Inventory ^43^. The experiment was approved by the Ethical Committee of the Leibniz Research Centre for Working Environment and Human Factors. All participants gave their written informed consent prior to the beginning of the experimental procedure. Please find additional information in the methods section in the supplementary material.

### Procedure

In each trial, participants performed both a number classification and a working memory task. These tasks were framed by a relevance cue and a feedback screen, both supporting participants to adapt their mental effort in a task-specific way throughout the trial (see figure 1).

Each trial started with a relevance cue indicating on which of the two tasks participants should focus their attention in the following trial. The cue was a triangle with either the number five in it (indicating that the number classification task should be focused) or a triangle with vertical stripes indicating that the working memory task should be focused (see figure 1). The relevance cue was presented for 500 ms and was followed by a fixation dot for 2500 ms. After that, the working memory task started with two Gabor patches (memory items) appearing on the left and right side of the screen for 400 ms, followed by a fixation dot for 2200 ms. Participants were instructed to remember the orientation of both patches. During the retention interval of the working memory task, participants performed a cued number-classification task, in which they classified a digit (number classification task target) as either smaller or bigger than five or as odd or even by pressing one of two buttons. A letter (classification cue, A, B, X or Y) appearing on screen for 200 ms indicated which of two classifications needed to be done. Bigger/smaller five was assigned to A or B and odd/even to X or Y for half of the participants, for the other half it was the other way round. The following classification digit (1, 2, 3, 4, 6, 7, 8, or 9) was presented for 200 ms. Its presentation was framed by a fixation dot period of 1000 ms before its appearance and 2600 ms after its disappearance. Responses to the number classification task were registered in the time interval starting 200 ms and ending 1800 ms after the onset of the digit. Responses outside this time interval were regarded as missing answers. Then, the working memory task continued with a retro-cue indicating which of the two Gabor Patches would be asked for in the following working memory retrieval phase. The retro-cue was an arrow, presented for 200 ms, pointing to the side (left or right) on which the to be retrieved Gabor patch had been presented before. It was followed by a fixation dot for 1000 ms. After that, the working memory target, a Gabor Patch with a random orientation, appeared on the screen. Participants were instructed to rotate the orientation of this Gabor patch in the same orientation as the retro-cued memory item using a computer mouse. If no response was made within the time interval of 200 ms to 4500 ms after the appearance of the target stimulus, the trial continued and the working memory answer was registered as missing. After a fixation dot interval of 200 ms, the trial ended with a feedback score appearing on the screen for 300 ms. The feedback score was calculated based on performance in both tasks, however the performance in the cued, i.e., in the more important, task was weighted three times as much as the performance in the un-cued task (for details see below). The ITI was 1200 ms.

After arrival, subjects were informed in detail about the experimental procedure. They were instructed to always perform both tasks but to put more effort in the respectively cued task than in the un-cued task. To motivate them to follow the task instructions, participants were additionally told that in case they collected more feedback points than 50 % of all participants, they had the opportunity to take part in a lottery. In fact, the lottery was done including all participants to not disadvantage less skillful subjects. Three participants won money in the lottery (€50, €45 and €30, respectively) which they got in addition to their normal study incentive. In addition, it was explained in detail how feedback scores were calculated.

Feedback points (FP) were assigned to responses in the following way: The maximal amount of feedback points (max) was 75 for the cued task and 25 for the un-cued task. In each trial, feedback points received for both tasks were added, and the sum (number between zero and 100) was depicted on screen accompanied by a percentage sign. Thus, performance in the cued task weighted three time as much as the performance in the un-cued task. In the number classification task, missing and incorrect answers always resulted in zero points. A response time of 200-400 ms with respect to the number classification digit resulted in the highest possible score (max). In the working memory task, missing answers or degree deviations between target and cued memory item bigger than 45° resulted in zero points. Apart from that, feedback points were assigned with the following formulas to reaction times (RT) or degree deviations:

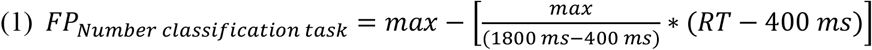

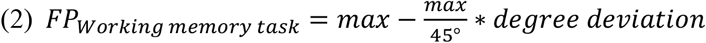

Thus, in the number classification task, subjects got a high amount of feedback points by answering as quickly and as accurately as possible. In the working memory task, they should answer as accurately as possible in order to be successful. In general, they had a higher likelihood to reach high scores if they varied their effort throughout the trial.

The experiment consisted of 10 blocks with 60 trials each, providing the possibility for breaks in between blocks. Trial order within each block was randomized. The levels of the experimental factors prioritized task (working memory task more important vs. number classification task more important), retro-cue orientation (left vs. right) and number classification type (odd / even vs. smaller / bigger five) appeared equally often within each block and co-occurrences were balanced. Participants needed roughly 2.5 hours for the 10 blocks. During this time, EEG was continuously recorded. Please find additional information regarding the experimental procedure and stimulus material in the supplementary material.

### EEG data acquisition and preprocessing

EEG was recorded using BrainVision Brainamp DC amplifier, BrainVision Recording software and 64 Ag/AgCl actiCAP slim active electrodes arranged according to the international 10-10 system (BrainProducts, Gilching, Germany). The ground electrode was placed at AFz and FCz served as online reference. Data was recorded with a sampling rate of 1000 Hz and impedances were kept below 10 kΩ during recording.

EEG preprocessing was based on EEGLAB ^44^ in combination with custom MATLAB code (R2020a, The MathWorks Inc., Natick, Massachusetts). Data were down-sampled to 200 Hz, high-pass filtered (FIR filter with Hamming window, order: 1321, transition band-width: 0.5 Hz, −6dB cutoff: 0.25 Hz, passband-edge: 0.5 Hz), low-pass filtered (FIR filter with Hamming window, order: 67, transition band-width: 10 Hz, −6dB cutoff: 45 Hz, passband-edge: 40 Hz) and re-referenced to CPz in order to be able to include the former reference position FPz in the analysis. Next, channels with poor data quality were detected and excluded from further analysis based on kurtosis and probability criteria (*M* = 0.54 channels, *SD* = 0.75 channels). After re-referencing to common average reference, data were segmented in epochs starting 3700 ms prior to and ending 13700 ms after the relevance cue. A baseline subtraction was done using the mean of each trial. Then, trials containing artifacts were detected and deleted automatically. On average, 10.71 % of trials were removed (*SD* = 7.92%). An independent component analysis (ICA) was performed on a random sample of the data (200 epochs). Prior to ICA, a dimensionality reduction was done, using a standard PCA and removing the 3 dimensions that explained least variance, to deal with potential linear dependencies amongst channels. After the ICA, on average, 31 % independent components (*SD* = 10.47%) were removed per participant using the EEGLab plugin ICLabel (Pion-Tonachini et al., 2019). Finally, previously removed channels were interpolated. Please find more details on EEG preprocessing in the method section of supplementary material.

### Time frequency analysis

Data were decomposed into frequency bands using complex Morlet wavelet convolution as described in Cohen ^46^. The frequencies of the wavelets ranged from 2-20 Hz (19 wavelets, linearly spaced). They had a full width of half maximum (fwhmf, Cohen, 2019) between 0.75 Hz for the lowest frequency and 4.25 Hz for the highest frequency, which corresponds to an fwhmf of 1000 to 200 ms in the time domain. Power values were extracted from the complex convolution result. To diminsih edge artifacts, the first and the last 700 ms of each epoch was deleted after the convolution. Raw power values were decibel-normalized using condition-unspecific baseline values based on the mean of the trials belonging to both conditions in the time interval from 700 to 200 ms prior to the relevance cue.

Lateralized power was computed based on raw power by pairing each electrode on the left hemisphere with its corresponding channel on the right hemisphere. For each electrode pair, we calculated one power average over all trials for contralateral electrodes (i.e., right hemisphere electrodes on trials in which the working memory task cue pointed to the left and left hemisphere electrodes on trials in which the working memory task cue pointed to the right) and one average for ipsilateral electrodes. Trials used for analysis were randomly drawn in a way that each combination of retro-cue direction (left vs. right) and response side in number classification (left vs. right) occurred equally often. To get a normalized asymmetry measure, we then computed a lateralization index ^48–51^ as follows:

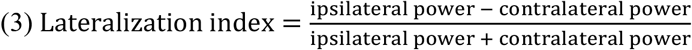

The index was calculated for every electrode pair, frequency, time point and subject, separately. Based on previous literature ^29,30,52^, we were interested in posterior alpha asymmetry which has been observed to be a correlate of attentional shifts within working memory. We therefore averaged the index values across a posterior electrode cluster (T7/T8, C5/C6, C3(C4, C1/C2, TP9/TP10, TP7/TP8, CP5/CP6, CP3/CP4, CP1/CP2, P7/P8, P5/P6, P3/4, P1/P2, PO7/PO8, PO3/PO4, PO9/PO10, O1/O2) and across frequencies ranging from 8 to 15 Hz (see figure 3).

### Decoding analysis

We did separate decoding analyses for both experimental conditions, i.e. for trials in which the number classification task was more important and trials in which the working memory task was more important. Within these two conditions, we decoded from the scalp topography the trial’s number classification type (smaller/bigger five vs. odd /even).

Decoding was done both based on broadband ERP data as well as based on alpha power. The decoding procedure was the same in both analyses except for the features used for training the models: For ERP decoding, the preprocessed EEG data of all channels were used as features, whereas for alpha power decoding time-frequency decomposed data at eight, nine, 10, 11 and 12 Hz of all channels were used. Employing Matlab scripts provided by Bae & Luck ^53^, we calculated classification accuracies based on a cross-validation with linear support vector machines, applied to trial averages rather than to single trials to boost signal-to-noise ratio. See supplementary material for more details.

### Statistical analysis

For details regarding statistical analysis of behavioral data see supplementary material. To statistically analyze EEG data, cluster-based permutation tests ^54^ were applied as they provide an elegant way to correct for multiple comparisons in high-dimensional EEG data.

In the non-lateralized time-frequency data analysis, the two experimental conditions (number classification task important and working memory task important) were compared at each pixel in the electrode x time point x frequency space to exploratively identify time points, frequencies and electrodes at which effort manipulation affects EEG activity. To this end, a cluster-based permutation test was conducted on decibel-normalized power values using the Matlab toolbox Fieldtrip ^55^. The test was computed across all frequencies obtained from time-frequency decomposition, i.e. across 2-20 Hz, across all 64 electrodes and across the whole epoch starting 3000 ms prior to and ending 13000 ms after the relevance cue (see also supplementary material).

Statistical analysis of decoding accuracies and power asymmetry values was done in two different ways: exploratively, using cluster-based permutation tests and hypothesis-driven, with Bayesian analyses applied to time intervals of interest. The cluster-based permutation tests were run over all time points of the trial between the relevance cue and 1000 ms after the onset of the working memory task target display. They were computed with the two Matlab functions *cluster_test*() and *cluster_test_helper*() which are part of the online code provided by Wolff et al. ^56^. See supplementary material for more details regarding these tests.

To test for general effects, cluster-based permutation tests were used to compare lateralization index against the case of zero asymmetry and decoding accuracies against chance level. To test for effects of the experimental manipulation, decoding accuracies and lateralization indices of working memory task important trials were compared against the respective measure obtained from number classification task important trials. To facilitate result interpretation, a second approach was applied for decoding and lateralization index analysis: a time interval of interest analysis including Bayesian statistics. In frequentist statistics, higher p-values do not indicate more evidence for the null hypothesis. In contrast, Bayesian statistics quantifies the evidence in favor for one of both hypotheses (alternative hypothesis H1 and null hypothesis H0). The Bayes factor B01 is the likelihood ratio expressing this evidence. More exactly, B_01_ is the likelihood for the measured data given that H0 is true divided by the likelihood for the measured data given that H1 is true. Thus, B_01_=2 means that the data are twice as likely H0 than under H1. Keysers & Wagenmakers ^57^ suggest that whereas 1/3<B_01_<3 provides not enough evidence for either hypothesis, B_01_>3 indicates substantial evidence for H0 and likewise, B_10_<1/3 indicates substantial evidence for H1 (see also Jeffreys, ^58^). A B_01_>10 is considered as strong evidence for H0. The time interval of interest analysis comprised two steps: We first calculated t-tests using Matlab. We then computed Bayes factors based on the outputs derived from the frequentist t-test as suggested by Morey & Wagenmaker ^59^, using a Matlab toolbox from Bart Krekelberg ^60^. Default effect size priors from the toolbox were used (Cauchy distribution scale = 0.707). Using this approach, we compared on the one hand lateralization indices versus zero / decoding accuracies versus chance levels and on the other hand values in number classification task important trials versus values in working memory task important trials. Two-sided one-sample *t* tests were computed in the first case and two-sided paired *t* tests in the other case.

Effect sizes were calculated for significant effects. η refers to adjusted partial eta squared ^61^. This is a bias-free effect size based on variance which is suited to show were pixels of a cluster are most pronounced. D refers to Cohen’s d. In some cases, both effect sizes are reported to provide a better comparability with other research.

## Data / code availability statement

Data and analysis code related to this manuscript will be made available online as an OSF repository upon publication.

## Acknowledgements

We would like to thank Pia Deltenre, Svenja Pschichholz, Aslisan Yavuz, Vivien Szczepanski, Joel Althoff, and Karin Lukaszewski for their assistance in collecting the data as well as Tobias Blanke for programming the experiment.

## Additional Information

Portions of these findings were presented as a poster at the 2nd workshop on mental effort, virtual conference, 2021. We have no conflicts of interest to disclose.

## Author contributions

N.L..: Conceptualization; Data collection; Formal analysis; Methodology; Visualization; Writing-original draft;Writing-review& editing. D.S.: Conceptualization; Writing-review& editing; Supervision. E.W.: Writing-review& editing; Supervision. S.A.: Conceptualization; Formal analysis; Methodology; Writing-original draft;Writing-review& editing; Supervision

## References

1. Shenhav, A. et al. Toward a Rational and Mechanistic Account of Mental Effort. Annu. Rev. Neurosci. 40, 99–124 (2017).

2. Inzlicht, M., Shenhav, A. & Olivola, C. Y. The Effort Paradox: Effort Is Both Costly and Valued. Trends in Cognitive Sciences 22, 337–349 (2018).

3. Shenhav, A., Botvinick, M. M. & Cohen, J. D. The Expected Value of Control: An Integrative Theory of Anterior Cingulate Cortex Function. Neuron 79, 217–240 (2013).

4. Botvinick, M. & Braver, T. Motivation and Cognitive Control: From Behavior to Neural Mechanism. Annual Review of Psychology 66, 83–113 (2015).

5. Yee, D. M. & Braver, T. S. Interactions of Motivation and Cognitive Control. Curr Opin Behav Sci 19, 83–90 (2018).

6. Failing, M. & Theeuwes, J. Selection history: How reward modulates selectivity of visual attention. Psychon Bull Rev 25, 514–538 (2018).

7. Krebs, R. M. & Woldorff, M. G. Cognitive Control and Reward. in The Wiley Handbook of Cognitive Control 422–439 (John Wiley & Sons, Ltd, 2017). doi:10.1002/9781118920497.ch24.

8. Buschman, T. J. & Kastner, S. From Behavior to Neural Dynamics: An Integrated Theory of Attention. Neuron 88, 127–144 (2015).

9. Miller, E. K. & Cohen, J. D. An Integrative Theory of Prefrontal Cortex Function. Annual Review of Neuroscience 24, 167–202 (2001).

10. Knutson, B., Westdorp, A., Kaiser, E. & Hommer, D. FMRI Visualization of Brain Activity during a Monetary Incentive Delay Task. NeuroImage 12, 20–27 (2000).

11. Schevernels, H., Krebs, R. M., Santens, P., Woldorff, M. G. & Boehler, C. N. Task preparation processes related to reward prediction precede those related to task-difficulty expectation. NeuroImage 84, 639–647 (2014).

12. van den Berg, B., Krebs, R. M., Lorist, M. M. & Woldorff, M. G. Utilization of reward-prospect enhances preparatory attention and reduces stimulus conflict. Cogn Affect Behav Neurosci 14, 561–577 (2014).

13. Sawaki, R., Luck, S. J. & Raymond, J. E. How Attention Changes in Response to Incentives. J Cogn Neurosci 27, 2229–2239 (2015).

14. Frömer, R., Lin, H., Dean Wolf, C. K., Inzlicht, M. & Shenhav, A. Expectations of reward and efficacy guide cognitive control allocation. Nat Commun 12, 1030 (2021).

15. Braver, T. S. The variable nature of cognitive control: a dual mechanisms framework. Trends Cogn.Sci. (Regul. Ed.) 16, 106–113 (2012).

16. Chiew, K. S. & Braver, T. S. Temporal Dynamics of Motivation-Cognitive Control Interactions Revealed by High-Resolution Pupillometry. Front Psychol 4, 15 (2013).

17. Chiew, K. S. & Braver, T. S. Dissociable influences of reward motivation and positive emotion on cognitive control. Cogn Affect Behav Neurosci 14, 509–529 (2014).

18. Fröber, K. & Dreisbach, G. The differential influences of positive affect, random reward, and performance-contingent reward on cognitive control. Cogn Affect Behav Neurosci 14, 530–547 (2014).

19. Fröber, K. & Dreisbach, G. How performance (non-)contingent reward modulates cognitive control. Acta Psychol (Amst) 168, 65–77 (2016).

20. Locke, H. S. & Braver, T. S. Motivational influences on cognitive control: behavior, brain activation, and individual differences. Cogn Affect Behav Neurosci 8, 99–112 (2008).

21. Fischer, R., Fröber, K. & Dreisbach, G. Shielding and relaxation in multitasking: Prospect of reward counteracts relaxation of task shielding in multitasking. Acta Psychol (Amst) 191, 112–123 (2018).

22. Gopher, D., Brickner, M. & Navon, D. Different difficulty manipulations interact differently with task emphasis: Evidence for multiple resources. Journal of Experimental Psychology: Human Perception and Performance 8, 146–157 (1982).

23. Gopher, D., Weil, M. & Siegel, D. Practice under changing priorities: An approach to the training of complex skills. Acta Psychologica 71, 147–177 (1989).

24. Lehle, C. & Hübner, R. Strategic capacity sharing between two tasks: evidence from tasks with the same and with different task sets. Psychological Research 73, 707 (2008).

25. Grootswagers, T., Wardle, S. G. & Carlson, T. A. Decoding Dynamic Brain Patterns from Evoked Responses: A Tutorial on Multivariate Pattern Analysis Applied to Time Series Neuroimaging Data. J Cogn Neurosci 29, 677–697 (2017).

26. Sharifian, F., Schneider, D., Arnau, S. & Wascher, E. Decoding of cognitive processes involved in the continuous performance task. International Journal of Psychophysiology 167, 57–68 (2021).

27. Etzel, J. A., Cole, M. W., Zacks, J. M., Kay, K. N. & Braver, T. S. Reward Motivation Enhances Task Coding in Frontoparietal Cortex. Cereb Cortex 26, 1647–1659 (2016).

28. Hall-McMaster, S., Muhle-Karbe, P. S., Myers, N. E. & Stokes, M. G. Reward Boosts Neural Coding of Task Rules to Optimize Cognitive Flexibility. J. Neurosci. 39, 8549–8561 (2019).

29. Poch, C., Campo, P. & Barnes, G. R. Modulation of alpha and gamma oscillations related to retrospectively orienting attention within working memory. Eur J Neurosci 40, 2399–2405 (2014).

30. Schneider, D., Mertes, C. & Wascher, E. On the fate of non-cued mental representations in visuo-spatial working memory: Evidence by a retro-cuing paradigm. Behavioural Brain Research 293, 114–124 (2015).

31. Kiss, M., Driver, J. & Eimer, M. Reward priority of visual target singletons modulates event-related potential signatures of attentional selection. Psychol Sci 20, 245–251 (2009).

32. Schneider, D., Bonmassar, C. & Hickey, C. Motivation and short-term memory in visual search: Attention’s accelerator revisited. Cortex 102, 45–56 (2018).

33. Beck, S. M., Locke, H. S., Savine, A. C., Jimura, K. & Braver, T. S. Primary and Secondary Rewards Differentially Modulate Neural Activity Dynamics during Working Memory. PLOS ONE 5, e9251 (2010).

34. Engelmann, J. B., Damaraju, E., Padmala, S. & Pessoa, L. Combined effects of attention and motivation on visual task performance: Transient and sustained motivational effects. Frontiers in Human Neuroscience 3, (2009).

35. Kostandyan, M. et al. Are all behavioral reward benefits created equally? An EEG-fMRI study. NeuroImage 215, 116829 (2020).

36. Cavanagh, J. F. & Frank, M. J. Frontal theta as a mechanism for cognitive control. Trends Cogn.Sci. (Regul. Ed.) 18, 414–421 (2014).

37. de Vries, I. E. J., Slagter, H. A. & Olivers, C. N. L. Oscillatory Control over Representational States in Working Memory. Trends in Cognitive Sciences 24, 150–162 (2020).

38. Klimesch, W. EEG alpha and theta oscillations reflect cognitive and memory performance: a review and analysis. Brain Research Reviews 29, 169–195 (1999).

39. Hefer, C. & Dreisbach, G. The motivational modulation of proactive control in a modified version of the AX-continuous performance task: Evidence from cue-based and prime-based preparation. Motivation Science 2, 116–134 (2016).

40. Krebs, R. M., Boehler, C. N., Roberts, K. C., Song, A. W. & Woldorff, M. G. The involvement of the dopaminergic midbrain and cortico-striatal-thalamic circuits in the integration of reward prospect and attentional task demands. Cereb Cortex 22, 607–615 (2012).

41. Padmala, S. & Pessoa, L. Reward reduces conflict by enhancing attentional control and biasing visual cortical processing. J Cogn Neurosci 23, 3419–3432 (2011).

42. Small, D. M. et al. Monetary incentives enhance processing in brain regions mediating top-down control of attention. Cereb Cortex 15, 1855–1865 (2005).

43. Oldfield, R. C. The assessment and analysis of handedness: The Edinburgh inventory. Neuropsychologia 9, 97–113 (1971).

44. Delorme, A. & Makeig, S. EEGLAB: an open source toolbox for analysis of single-trial EEG dynamics including independent component analysis. J Neurosci Methods 134, 9–21 (2004).

45. Pion-Tonachini, L., Kreutz-Delgado, K. & Makeig, S. ICLabel: An automated electroencephalographic independent component classifier, dataset, and website. Neuroimage 198, 181–197 (2019).

46. Cohen, M. X. Chapter 13: Complex Morlet Wavelets and Extracting Power and Phase. in Analyzing neural time series data: theory and practice 151–174 (The MIT Press, 2014).

47. Cohen, M. X. A better way to define and describe Morlet wavelets for time-frequency analysis. NeuroImage 199, 81–86 (2019).

48. Haegens, S., Händel, B. F. & Jensen, O. Top-down controlled alpha band activity in somatosensory areas determines behavioral performance in a discrimination task. J Neurosci 31, 5197–5204 (2011).

49. Klatt, L.-I., Getzmann, S., Begau, A. & Schneider, D. A dual mechanism underlying retroactive shifts of auditory spatial attention: dissociating target- and distractor-related modulations of alpha lateralization. Sci Rep 10, 13860 (2020).

50. Wildegger, T., van Ede, F., Woolrich, M., Gillebert, C. R. & Nobre, A. C. Preparatory α-band oscillations reflect spatial gating independently of predictions regarding target identity. J Neurophysiol 117, 1385–1394 (2017).

51. Wöstmann, M., Herrmann, B., Maess, B. & Obleser, J. Spatiotemporal dynamics of auditory attention synchronize with speech. Proceedings of the National Academy of Sciences 113, 3873–3878 (2016).

52. Rösner, M., Arnau, S., Skiba, I., Wascher, E. & Schneider, D. The spatial orienting of the focus of attention in working memory makes use of inhibition: Evidence by hemispheric asymmetries in posterior alpha oscillations. Neuropsychologia 142, 107442 (2020).

53. Bae, G.-Y. & Luck, S. J. Dissociable Decoding of Spatial Attention and Working Memory from EEG Oscillations and Sustained Potentials. J Neurosci 38, 409–422 (2018).

54. Maris, E. & Oostenveld, R. Nonparametric statistical testing of EEG- and MEG-data. J Neurosci Methods 164, 177–190 (2007).

55. Oostenveld, R., Fries, P., Maris, E. & Schoffelen, J.-M. FieldTrip: Open source software for advanced analysis of MEG, EEG, and invasive electrophysiological data. Comput Intell Neurosci 2011, 156869 (2011).

56. Wolff, M. J., Jochim, J., Akyürek, E. G. & Stokes, M. G. Dynamic hidden states underlying working-memory-guided behavior. Nat. Neurosci. 20, 864–871 (2017).

57. Keysers, C., Gazzola, V. & Wagenmakers, E.-J. Using Bayes factor hypothesis testing in neuroscience to establish evidence of absence. Nat Neurosci 23, 788–799 (2020).

58. Jeffreys, H. Theory of probability. (Clarendon Press; Oxford University Press, 1998).

59. Morey, R. D. & Wagenmakers, E.-J. Simple relation between Bayesian order-restricted and point-null hypothesis tests. Statistics & Probability Letters 92, 121–124 (2014).

60. Krekelberg, B. BayesFactor. GitHub https://github.com/klabhub/bayesFactor (2021).

61. Mordkoff, J. T. A simple method for removing bias from a popular measure of standardized effect size: Adjusted partial eta squared. Advances in Methods and Practices in Psychological Science 2, 228–232 (2019).

62. Botvinick, M., Huffstetler, S. & McGuire, J. T. Effort discounting in human nucleus accumbens. Cogn Affect Behav Neurosci 9, 16–27 (2009).

63. Cooper, P. S., Wong, A. S. W., McKewen, M., Michie, P. T. & Karayanidis, F. Frontoparietal theta oscillations during proactive control are associated with goal-updating and reduced behavioral variability. Biological Psychology 129, 253–264 (2017).

64. Arnau, S., Wascher, E. & Küper, K. Age-related differences in reallocating cognitive resources when dealing with interruptions. Neuroimage 191, 292–302 (2019).

65. Min, B.-K. & Park, H.-J. Task-related modulation of anterior theta and posterior alpha EEG reflects top-down preparation. BMC Neuroscience 11, 79 (2010).

66. Bunzeck, N., Guitart-Masip, M., Dolan, R. J. & Düzel, E. Contextual novelty modulates the neural dynamics of reward anticipation. J Neurosci 31, 12816–12822 (2011).

67. Doñamayor, N., Schoenfeld, M. A. & Münte, T. F. Magneto- and electroencephalographic manifestations of reward anticipation and delivery. NeuroImage 62, 17–29 (2012).

68. Sauseng, P., Griesmayr, B., Freunberger, R. & Klimesch, W. Control mechanisms in working memory: A possible function of EEG theta oscillations. Neuroscience & Biobehavioral Reviews 34, 1015–1022 (2010).

69. Kawasaki, M. & Yamaguchi, Y. Frontal theta and beta synchronizations for monetary reward increase visual working memory capacity. Soc Cogn Affect Neurosci 8, 523–530 (2013).

70. Cunillera, T. et al. Brain oscillatory activity associated with task switching and feedback processing. Cogn Affect Behav Neurosci 12, 16–33 (2012).

71. Rawle, C., Miall, C. & Praamstra, P. Frontoparietal theta activity supports behavioral decisions in movement-target selection. Frontiers in Human Neuroscience 6, 138 (2012).

72. van Driel, J., Swart, J. C., Egner, T., Ridderinkhof, K. R. & Cohen, M. X. (No) time for control: Frontal theta dynamics reveal the cost of temporally guided conflict anticipation. Cogn Affect Behav Neurosci 15, 787–807 (2015).

73. Allen, J. J. B., Coan, J. A. & Nazarian, M. Issues and assumptions on the road from raw signals to metrics of frontal EEG asymmetry in emotion. Biol Psychol 67, 183–218 (2004).

74. Iemi, L., Chaumon, M., Crouzet, S. M. & Busch, N. A. Spontaneous Neural Oscillations Bias Perception by Modulating Baseline Excitability. J. Neurosci. 37, 807–819 (2017).

75. Jensen, O. & Mazaheri, A. Shaping Functional Architecture by Oscillatory Alpha Activity: Gating by Inhibition. Front. Hum. Neurosci. 4, (2010).

76. Schneider, D., Herbst, S. K., Klatt, L.-I. & Wöstmann, M. Target enhancement or distractor suppression? Functionally distinct alpha oscillations form the basis of attention. Eur J Neurosci (2021) doi:10.1111/ejn.15309.

77. Bartsch, F., Hamuni, G., Miskovic, V., Lang, P. J. & Keil, A. Oscillatory brain activity in the alpha range is modulated by the content of word-prompted mental imagery. Psychophysiology 52, 727–735 (2015).

78. Bazanova, O. M. & Vernon, D. Interpreting EEG alpha activity. Neurosci Biobehav Rev 44, 94–110 (2014).

79. Wang, C., Rajagovindan, R., Han, S.-M. & Ding, M. Top-Down Control of Visual Alpha Oscillations: Sources of Control Signals and Their Mechanisms of Action. Front Hum Neurosci 10, 15 (2016).

80. Fischer, R. & Plessow, F. Efficient multitasking: parallel versus serial processing of multiple tasks. Front Psychol 6, (2015).

81. Ewing, K. C. & Fairclough, S. H. The Effect of an Extrinsic Incentive on Psychophysiological Measures of Mental Effort and Motivational Disposition when Task Demand is Varied. Proceedings of the Human Factors and Ergonomics Society Annual Meeting 54, 259–263 (2010).

82. Klink, P. C., Jeurissen, D., Theeuwes, J., Denys, D. & Roelfsema, P. R. Working memory accuracy for multiple targets is driven by reward expectation and stimulus contrast with different time-courses. Sci Rep 7, 9082 (2017).

83. Leon, M. I. & Shadlen, M. N. Effect of Expected Reward Magnitude on the Response of Neurons in the Dorsolateral Prefrontal Cortex of the Macaque. Neuron 24, 415–425 (1999).

84. Grogan, J. P., Randhawa, G., Kim, M. & Manohar, S. G. Motivation Improves Working Memory by Two Processes: Prioritisation and Retrieval Thresholds. (2021) doi:10.31234/osf.io/qkycj.

