## Supplementary material for "Task prioritization modulates low frequency EEG dynamics reflecting proactive cognitive control"

**[1]Leibniz Research Centre for Working Environment and Human Factors, Dortmund, Germany**

### Supplementary methods

#### Participants

Thirty-three volunteers participated in the experiment. One dataset was excluded from analysis due to bad data quality; four additional datasets were excluded due to an error in the stimulus presentation program. The remaining twenty-eight subjects were included in the analysis. Participants received course credit, or they were paid €10 per hour. They also had the opportunity to win an additional amount of money by participating in a lottery.

#### Stimulus material

Stimuli were presented using FreePascal software (https://www.freepascal.org/) on a 32 in., 1920 × 1080 pixels VSG monitor (Display++ LCD) with 100 Hz refreshing rate. Throughout the experiment, the screen background color was grey (CIE1931: 0.287, 0.312, 10). All stimuli were presented at the center of the screen with a viewing distance of 1450 cm, except for the two memory items which had their center at a visual angle of 1.5° on the left and right side of the screen's central point. Letters (height: 1°, font type: Arial), digits (height: 1°, font type: Arial), feedback scores (height: 1°, font type: Arial), retro-cues (width: 1°), relevance cues (height: 2.8°) and fixation dots (diameter: 0.18°) were presented in white (CIE1931: 0.287, 0.312, 50). The Gabor patches (diameter: 2°) had a contrast of 85%, a spatial frequency of 3.25 cycles per degree, and a phase of 180°. One of 180 possible orientations (1° to 180° in 1° steps) was randomly assigned to each Gabor patch (target, cued and un-cued item) in each trial, with the only limitation that cued and un-cued item's orientations differed with more than 15°.

#### Procedure and post-EEG questionnaire

After the electrodes were montaged, subjects performed 40-120 practice trials prior to the start of the experiment. The practice phase ended when error rates and response times remained stable.

During the experiment, participants sat in a comfortable armchair handling a response mouse with their right hand. The buttons of the response mouse were soldered to a custom response device with very low latency (<1ms, no jitter) while leaving the mouse cursor, i.e., the optical sensor of the mouse, operational. In the NC task, participants used their index finger to click left to give the answer "smaller five" or "odd" and their ring finger to click right to give the answer "larger than five" or "even". In the working memory task, participants rotated the orientation of the Gabor patch by laterally shifting the mouse. A left click on the mouse locked the final orientation in place.

At the end of the experiment, participants completed a short paper-based questionnaire (post-EEG questionnaire). They gave their answers by marking a spot on a continuous bar of 10 cm length which was labelled at zero cm (left label) and 10 cm (right label). In specific, they were asked the following questions: “How demanding were the number tasks for you?“, left label: not at all demanding, right label: very demanding. “How motivated were you to do the number tasks?“, left label: not at all motivated, right label: very motivated. “Please indicate in how far you agree to the following statement. I tried harder in the number task, when it was important than when it was unimportant.”, left label: I agree not at all, right label: I totally agree. The same three questions were asked for the working memory task (“number task” replaced by “memory task”).

**EEG preprocessing**

To detect and remove artifactual channels, the EEGLab function *pop_rejchan*() was used. A kurtosis value per channel based on all data points of that channel was calculated. Channels with z-transformed (over all channels) kurtosis values lower than -10 or higher than 10 were rejected. Z-transformation was done based on standard deviation and mean of a trimmed (10% highest values and 10% lowest values removed) kurtosis distribution. Likewise, joint probability values, that is the negative sum of logarithmized density values of z-transformed data points of the channel, were calculated, z-transformed (based on a trimmed distribution) and channels with a lower z-transformed joint probability than minus five or a higher z-transformed joint probability than five were removed.

Trials with artifacts were automatically deleted using *pop_autorej*(). This EEGlab function rejected trials containing fluctuation larger than 1000 µV and then iteratively detected further unregular epochs based on single trials’ joint probability values, whereas maximally five % of trials were rejected per iteration. The probability threshold was set to five, meaning that z-transformed joint probability values that are smaller than minus five or higher than five are detected. Z-transformation of joint probability values was based on mean and standard deviation of a 20 % trimmed joint probability distribution over epochs, calculated for every channel.

ICA was done based on the function *runica*(). The function *iclabel*() was applied on obtained ICs. This EEGLab plugin automatically categorizes ICs based on a database of manually categorized data. ICs that represented an artifact (eye movements /blinks, pulse or muscle artifacts, line noise, channel noise) with more than 50 % likelihood according to this categorization were removed.

Channel were interpolated using the function *pop_interp*() with default parameters.

**Decoding analysis**

In detail, we conducted decoding analyses as follows: At the beginning, data were down-sampled to 20 Hz to improve processing speed. Then, at each time point, a 15-fold cross validation was performed 10 times: Each cross validation started with randomly assigning trials belonging to a class (i.e., to either smaller/bigger 5 or to odd/even) in 15 groups resulting in six to nine trials per group ($M$ = 8.05, $SD$ = 0.90). Group size was constant within each analysis and participant. If the number of trials belonging to each class was not divisible by 15, surplus trials were randomly determined and excluded from the analysis. A support vector machine was then trained on the 28 data points obtained after averaging across trials belonging to a group. Each data points had thus 64 dimensions (or 64 * 5 dimensions in alpha decoding) as values from all 64 electrodes (and five frequencies) were used. Training was done with the Matlab function *fitcecoc*(). In the testing phase, the remaining two data points, consisting of electrode or electrode / frequency values of the two (one per class) left-out group averages, were classified based on the output model created by *fitcecoc*() using the Matlab function *predict*(). Test- and training procedure was then repeated with each group being the left-out data point once, i.e., 15 times. Thus, at each time point, 150 (10 iterations x 15 groups) class predictions were done which could be correct (the class the group really belongs to was predicted) or incorrect (the other class was predicted). Decoding accuracy at each time point was determined as the number of correct predictions divided by 150. This results in a time series of decoding accuracies per participant. Finally, these time series were smoothed using a 5-point (+/- 100 ms) moving average window and used for statistical analyses.

**Statistical analysis**

Matlab (R2020a, The MathWorks Inc., Natick, Massachusetts) was used to perform statistical analyses.

**Statistical analysis of behavioral data**

In the working memory task, on average 0.20% of the number classification task important trials and 0.31% of working memory task important trials were not or not quickly enough answered. For the remaining trials, median degree deviation between the to-be-remembered and the answer orientation and median mouse onset times were calculated for each participant.

Median instead of mean over trials was chosen as the median is more robust against outliers and more reliable for most unimodal skewed distributions with continuous variables [1].

In the number classification task, on average 2.33% of responses in working memory task important trials and 1.96% of responses in the number classification task important trials were missing. Median RT across trials based on correct responses was calculated.

In addition, for each participant we computed mean feedback points.

Two-sided paired t-tests between the number classification task important and working memory task important trials were computed on resulting single subject accuracy, reaction time and feedback point data.

Furthermore, with each of the three question couples in the post-EEG questionnaire, a paired t-test was calculated on post-EEG questionnaire response scores (distance of marked position from the left side of the answer scale) between the scores referring to the number classification task question and the scores referring to the working memory task question, respectively. For one subject, questionnaire data was missing.

**Statistical analysis of time-frequency decomposed data**

Non-lateralized time-frequency data was statistically analyzed using the Matlab toolbox Fieldtrip. A test statistic for the permutation test was calculated based on the following procedure: Repeated measures two-sided *t*-tests at each data pixel in the electrode × time × frequency space were computed and above-threshold neighbored pixels identified as cluster. Pixels with a *p*-value under 0.05 in the *t*-test were considered above-threshold. The test statistic was the sum of the *t*-values of the pixels belonging to each cluster.

In order to identify significant clusters, a non-parametrical distribution of this test statistic was created under the H0 assumption that both experimental conditions do not differ. To this end, a Montecarlo simulation with 1000 random draws was computed based on the real data but with trials randomly assigned to experimental conditions. Then, in each of the 1000 generated random data sets, clusters and its corresponding t sums were identified and the biggest *t* sum per data set was incorporated into the H0 distribution.

Real data clusters were considered significant if they had a *t* sum that was higher than the 97,5% or lower than the 2,5% quantile of the H0 distribution.

**Statistical analysis of decoding accuracies and lateralized time-frequency decomposed data**

Decoding accuracies and lateralization indices were statistically compared using cluster-based permutation tests. Tests were calculated based on difference values for the two experimental conditions of interest (data vs. chance level; data vs. zero; working memory task important vs. number classification important). The test statistic was the sum of the means calculated at each pixel that belongs to a cluster. 10000 random data sets were created by randomly assigning a sign (plus or minus) to each data point of the real data (difference values between both experimental condition) and the cluster with the biggest mean sum was incorporated into the H0 distribution of mean sums. The real data clusters were considered significant if their mean sum was higher than the 97.5 % or lower than the 2.5 % quantile of the Ho distribution. Mean sums were computed as follows: Means of difference data over all subjects at every data point were computed. A permutation test revealed pixels with significant ($p<0.05$) differences regarding the two experimental conditions. These were considered above-threshold. Two neighbored above-threshold pixels or more add up to a cluster.

### Supplementary results

#### Post-EEG questionnaire and feedback points

Post-EEG questionnaire answers and feedback points revealed differences regarding perceived motivation and difficulty associated with both tasks:

The mean feedback score per participant was significantly higher in trials in which the number classification task was more important ($M$ = 74.80, $SD$ = 9.30) than in trials in which the working memory task was more important ($M$ = 68.45, $SD$ = 8.63; $t(27)=-9.57$, $p < 0.01$, $\eta = 0.76$, $d = 1.81$).

According to participants' self-reports, subjects were slightly but not significantly more motivated ($t(26)=-2.00$, $p = 0.06$) to do the number classification tasks ($M$ = 7.64, $SD$ = 2.09) than to do the working memory tasks ($M$ = 6.91, $SD$ = 2.16) and considered the working memory tasks ($M$ = 6.85, $SD$ = 1.62) significantly more demanding than the number classification tasks ($M$ = 4.65, $SD$ = 2.42; $t(26)=3.76$, $p < 0.01$, $\eta = 0.33$, $d = 0.72$).

These differences do not abolish the task-specific effort allocation throughout the trials that we were interested in in this study as performance data clearly shows. This is in line with participants self-reports regarding effort allocation. Their answers to the statement "I tried harder in the x task, when it was important than when it was unimportant." did not differ significantly ($t(26)=0.20$, $p = 0.84$) when x was the number classification ($M$ = 7.25, $SD$ = 2.04) and when x was the working memory ($M$ = 7.37, $SD$ = 2.48).

### References

1. von Hippel, P. T. Mean, Median, and Skew: Correcting a Textbook Rule. *Journal of Statistics Education* **13**, null (2005).
